# Somatic mutation phasing and haplotype extension using linked-reads in multiple myeloma

**DOI:** 10.1101/2024.08.09.607342

**Authors:** Steven M. Foltz, Yize Li, Lijun Yao, Nadezhda V. Terekhanova, Amila Weerasinghe, Qingsong Gao, Guanlan Dong, Moses Schindler, Song Cao, Hua Sun, Reyka G. Jayasinghe, Robert S. Fulton, Catrina C. Fronick, Justin King, Daniel R. Kohnen, Mark A. Fiala, Ken Chen, John F. DiPersio, Ravi Vij, Li Ding

## Abstract

Somatic mutation phasing informs our understanding of cancer-related events, like driver mutations. We generated linked-read whole genome sequencing data for 23 samples across disease stages from 14 multiple myeloma (MM) patients and systematically assigned somatic mutations to haplotypes using linked-reads. Here, we report the reconstructed cancer haplotypes and phase blocks from several MM samples and show how phase block length can be extended by integrating samples from the same individual. We also uncover phasing information in genes frequently mutated in MM, including *DIS3*, *HIST1H1E*, *KRAS*, *NRAS*, and *TP53*, phasing 79.4% of 20,705 high-confidence somatic mutations. In some cases, this enabled us to interpret clonal evolution models at higher resolution using pairs of phased somatic mutations. For example, our analysis of one patient suggested that two *NRAS* hotspot mutations occurred on the same haplotype but were independent events in different subclones. Given sufficient tumor purity and data quality, our framework illustrates how haplotype-aware analysis of somatic mutations in cancer can be beneficial for some cancer cases.

## Introduction

Human genomes are diploid with two copies of each autosomal chromosome. Homologous chromosomes are distinct because they represent unique patterns of germline variation inherited from each parent. While genotypes represent the alleles at a specific locus, haplotypes are defined as groups of alleles across many loci separated according to which homolog they come from. Variant phasing and haplotype reconstruction may be achieved through technological and computational methods with a variety of data types and integration strategies from large public databases and individual samples.^1–24^

Determining the haplotype of cancer-associated mutations informs our understanding of the oncogenic process, but that information is typically lost with next-generation bulk sequencing.^25,26^ Linked-read sequencing overcomes that limitation by labelling DNA from the same haplotype with the same barcode. Zheng *et al.* described this linked-read approach, accurately modeling fusion breakpoints and revealing biallelic *TP53* inactivation by phasing a mutation and hemizygous deletion to opposite haplotypes.^27^ Marks *et al.* established the accuracy and reliability of linked-reads and explored the impact of variant density and heterozygosity on phasing performance.^28^ Linked-reads have impacted cancer study design and are especially well-suited for structural variant detection.^29–38^ Greer, et al. compared gastric cancer metastases and delineated a complex structural variant leading to *FGFR2* amplification.^39^ Viswanathan, et al. determined the order of events in a cohort of prostate cancer patients, showing androgen receptor (*AR*) gene duplications and *CDK12* inactivation, phasing somatic mutations if the reads supporting it were assigned to a haplotype and phase block, and developing allele-specific copy number detection methods.^40,41^ Sereewattanawoot, et al. matched *cis*-acting regulatory variants with allele-specific expression in lung cancer cell lines.^42^ ENCODE cell lines K562 and HepG2 have been used for deeply-integrated linked-read investigations.^43,44^

In this study, we analyzed 23 samples from a cohort of 14 multiple myeloma patients using linked-read whole genome sequencing (lrWGS) generated using the 10X Genomics Chromium System. Multiple myeloma (MM) is the second most common form of blood cancer and has a median 5-year survival around 50%.^45^ MM is caused by clonal proliferation of plasma cells in the bone marrow. Primary genetic aberrations include hyperdiploidy and translocations that join the highly expressed IGH locus (chr14) with oncogenes, including t(11;14) (*CCND1*), t(4;14) (*WHSC1*), t(6;14) (*CCND3*), and t(14;20) (*MAFB*). Secondary events include *MYC* translocations and driver mutations. MAPK is the most commonly mutated pathway in MM, including somatic mutations in *KRAS*, *NRAS*, and *BRAF*.^45^ Better appreciation of the haplotype context of these events, both driver mutations and structural variations, is necessary to improve targeted therapies and understanding of myelomagenesis. We created a framework for systematically phasing somatic mutations to haplotypes, allowing for deeper interpretation of tumor evolution in some cases. We also illustrate the concept of extending phase blocks using shared germline information across samples from the same individual. Our cohort represents a large resource of multiple myeloma lrWGS data and improves our understanding of human haplotype and cancer haplotype analysis.

## Results

### Haplotype-aware methods build on phasing information to analyze somatic mutations

The advantage of lrWGS over traditional WGS is that reads mapping to the same genomic region with the same barcode most likely originated from the same piece of high molecular weight (HMW) DNA (Fig. 1a).^27^ The Long Ranger pipeline (10X Genomics) aligns reads, calls and phases variants, reports structural variants (SVs), and produces phasing quality metrics. With enough sequencing depth and allelic heterogeneity, Long Ranger is able to phase variants and reads. Variants and reads are grouped into phase blocks, defined as genomic ranges in which haplotype assignments are consistent. Within a phase block, all variants assigned to a certain haplotype are thought to have originated from the same biological haplotype. The haplotype order may switch in another phase block, so haplotype assignments cannot be compared between phase blocks. Long Ranger phasing is designed to work with germline variants and does not distinguish between germline variants and somatic mutations in cancer. Phasing performance may be suboptimal for somatic mutations with low variant allele frequency (VAF), in regions of copy number variation, and in tumor samples with low purity or heterogeneous clonal structure. Specific methods are necessary to overcome this limitation.^46^

**Figure 1.**
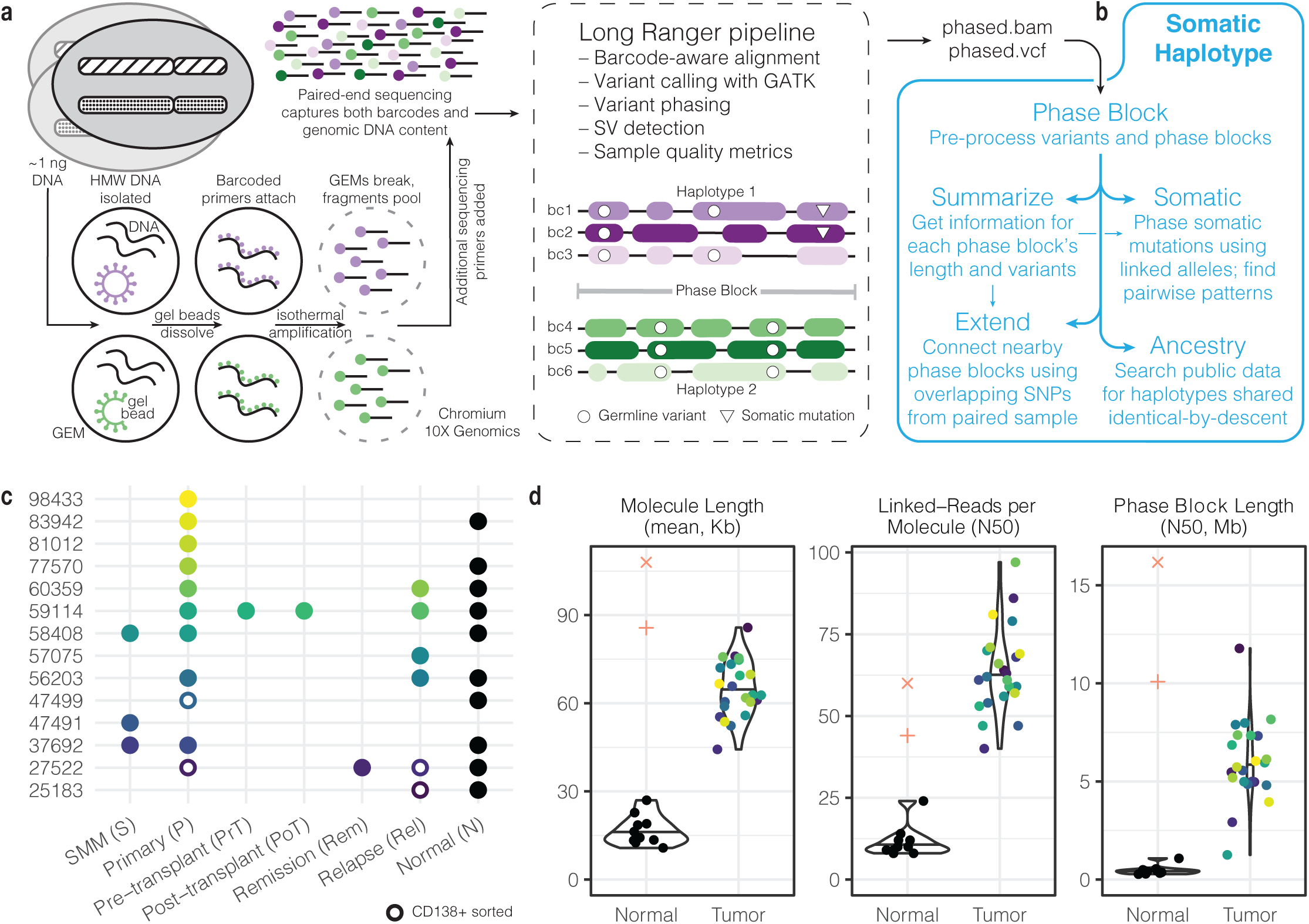
Linked-read data generation and analysis pipeline. **a.** The 10X Genomics Chromium platform tags large DNA molecules with barcodes such that reads originating from the same molecule have the same barcode. The Long Ranger pipeline aligns reads and phases variants. **b.** SomaticHaplotype builds upon Long Ranger output with several modules, including *phaseblock*, *summarize*, *somatic*, *extend*, and *ancestry*. **c.** Our cohort comprises 14 multiple myeloma patients across several disease stages for a total of 23 tumor samples. **d.** Quality control measures for our tumor and normal samples plus 1000 Genomes samples NA12878 (+) and NA19240 (x). Violin plots defined as: center line, median; violin limits, minimum and maximum values; points, every observation. Molecule Length (mean, Kb): length-weighted mean input DNA length in kilobases. Linked-Reads per Molecule (N50): N50 of read-pairs per input DNA molecule. Phase Block Length (N50, Mb): N50 length of phase blocks in megabases.

To enable further downstream processing of lrWGS data, we developed additional methods that use Long Ranger output to further analyze single nucleotide variant (SNV) mutations collectively referred to as SomaticHaplotype (Fig. 1b) (see **Methods** and **Code Availability**). Given the phased variant call format (VCF) file and phased bam file produced by Long Ranger, the *phaseblock* module constructs PhaseBlock and Variant objects with information derived from reads and variant calls for use by later modules. The *summarize* module reports summary information about each phase block, including genomic range and number of variants, and global statistics like phase block length N50. The *somatic* module uses two complementary approaches to assign high-confidence somatic mutations to haplotypes and then analyzes the haplotype relationship between proximal pairs of events. The *extend* module utilizes germline variation from matched samples to bridge gaps between phase blocks and suggests how to make neighboring phase blocks have consistent haplotype assignments. The *ancestry* module augments lrWGS data with information from large-scale phased resources, like the 1000 Genomes Project.

Our data set comprises lrWGS data from 14 patients diagnosed with multiple myeloma (Fig. 1c). Longitudinal samples were taken from the premalignant smoldering multiple myeloma (S), primary diagnosis (P), pre-transplant (PrT), post-transplant (PoT), remission (Rem), and relapse (Rel) stage. In total, 23 tumor samples and 10 skin normal samples were processed with lrWGS. Four tumor samples were CD138+ sorted to enrich for plasma cells, increasing tumor purity. Other samples were not CD138+ sorted and contain varying compositions of microenvironment cells along with tumor plasma cells. In addition, for 9 CD138+ sorted tumor samples with matched lrWGS, we generated whole genome sequencing (WGS) data with increased tumor purity to make high confidence somatic mutation calls (6 samples available at first data freeze) and structural variant calls (9 samples) (Supplementary Table 1; see **Methods**). Please see Supplementary Table 1 for tumor purity estimates of lrWGS samples with matched CD138+ sorted WGS samples (median tumor purity of sorted lrWGS = 0.676, n = 1; median tumor purity of unsorted lrWGS = .202, n = 4).

Cell-type composition, including tumor purity, shapes our interpretation of results from the cohort collectively and from individual samples. CD138+ sorting of four tumor samples selected for tumor-associated plasma cells, increasing tumor purity and our ability to detect interesting somatic mutation events. In unsorted samples comprising many immune and stromal cells not carrying the somatic mutations found in the tumor, we found tumor purity to be an important limiting factor that restricted our ability to more broadly generalize our findings across the dataset. Instead, we illustrate the types of analysis enabled by our framework by focusing on particular cases with the data quality sufficient for confident interpretation.

Quality control measures of our tumor samples compared well with data from publicly-available gold-standard data from two 1000 Genomes samples (see **Data Availability**) (Fig. 1d, Supplementary Figure 1, Supplementary Table 2). Molecule length refers to the size of the long, HMW DNA fragments. In our tumor samples, the mean molecule length per sample ranged from 44.3 Kb to 85.8 Kb with a median of 62.8 Kb, whereas in our normal skin samples, the median value was 15.3 Kb. Linked-reads per molecule is the number of read pairs originated from each molecule, and the N50 value indicates that half of the molecules have that many reads pairs or more. In our tumor samples, the N50 linked-reads per molecule ranged from 40 to 97 with a median of 62, compared to a median of 10 in our skin samples. Finally, the N50 phase block length in tumor samples ranged from 1.3 Mb to 11.8 Mb with a median of 5.7 Mb, whereas the median was 0.4 Mb in skin samples. Given the consistent lack of informative linked-read information in our skin samples, we excluded them from downstream analysis. The skin samples were only used as a control for somatic mutation calling from our sorted WGS samples. For tumor samples, the median corrected mass of input DNA loaded into the Chromium chip was 1.3 ng, and the median mean sequencing depth was 71.6 reads. The median percentage of single nucleotide variants (SNVs) phased by Long Ranger was 99.2%.

See Zhang, et al. for additional quality metrics that may be applied to linked-read data.^47^

### Phase block lengths reflect biologically-relevant genomic changes

We examined the distribution of phase block lengths to explore patterns in our data. N50 phase block lengths were consistent across chromosomes, with the median N50 ranging from 4.42 Mb on chr15 to 7.74 Mb on chr18 (Supplementary Figure 2a). Chr1 showed the least variation in N50 phase block length (median 4.52 Mb, standard deviation 1.37 Mb). Chr21 showed the greatest variation (median 5.78 Mb, standard deviation 9.33 Mb) and also had the highest overall values, with 6 samples having N50 phase block lengths above 20 Mb, 4 of which came from Patient 59114. Some samples, such as 25183 (Rel), had consistently higher N50 values across many chromosomes (Supplementary Figure 2b). This may be due to this sample having the highest mean molecule length (85.8 Kb) and percentage of mapped reads (97.7%) of all tumor samples. Another sample, 58408 (P), had consistently shorter phase blocks, but quality control measures did not clearly indicate why.

Chr13 and chr22 from 27522 (P) showed low N50 phase block lengths, and the distribution of phase block lengths from those two chromosomes is strikingly different from that of other chromosomes (Supplementary Figure 2c). The N50 phase block lengths for chr13 and chr22 were 0.42 Mb and 0.38 Mb, respectively, compared to that sample’s overall median N50 of 5.9 Mb. Both chr13 and chr22 had a one copy deletion across the entire chromosome, leading to a lack of heterozygosity needed for long phase blocks (Supplementary Figure 3). Hemizygous chr13 and chr22 phase blocks from 27522 (P) are much shorter across the entire chromosome compared to those from the remission sample, which is closer to an overall diploid state with low tumor content (Supplementary Figure 2d). However, we can interpret this sequencing artifact in a biologically meaningful way, and one benefit of homozygosity across an entire chromosome is the potential to resolve the entire chromosome’s haplotype structure. Deletion size, tumor purity, and the proportion of tumor cells with copy number loss are important factors determining the ability of deletion regions to be phased.

In total, phase blocks cover 60.6 Gb across our 23 tumor samples (Supplementary Figure 2e), for an average of 2.6 Gb per sample. 72.2% (32,426/44,918 phase blocks) of phase blocks are between 0 and 1 Mb, but those short segments account for only 8.4% (5.1/60.6 Gb) of the total amount of genome covered by phase blocks in these samples. In comparison, 3,776 phase blocks are between 1-2 Mb and cover 5.5 Gb (9.0%). The distribution of genomic coverage by phase blocks of increasing length has a right-skewed long tail distribution. There are 19 phase blocks longer than 30 Mb, and the longest phase block is 59.2 Mb. As expected, there is a strong linear relationship between phase block length and the number of phased heterozygous variants (r^2^ = 0.96). Over the 5.0 Mb human leukocyte antigen (HLA) region of chr6 (chr6:28510120-33480577), we observed a median of 4 phase blocks greater than 1kb in length (range 1-13 phase blocks), which covered between 93.5% and 100% of the region (median 98.7%). HLA region haplotyping could help match patients and donors before allogeneic stem cell transplants in limited and specific MM cases.^48^

### Somatic mutations can be phased to specific haplotypes using linked alleles

The haplotype context in which somatic mutations occur may be biologically relevant. For example, knowing the phase of two mutations affecting the same gene would indicate whether they cause biallelic inactivation or only alter one copy. However, tumor impurity, heterogeneity, and variable sequencing coverage make somatic mutations harder to identify and phase using standard approaches. To phase somatic mutations, we built upon the strengths of Long Ranger by examining germline variants that occur on each barcode associated with a somatic mutation site (Fig. 2a). We defined linked alleles as alleles co-occurring on the same barcode with either the reference or alternate allele at the somatic mutation site. We know that alleles co-occurring with the same barcode most likely originated from the same molecule of HMW

**Figure 2.**
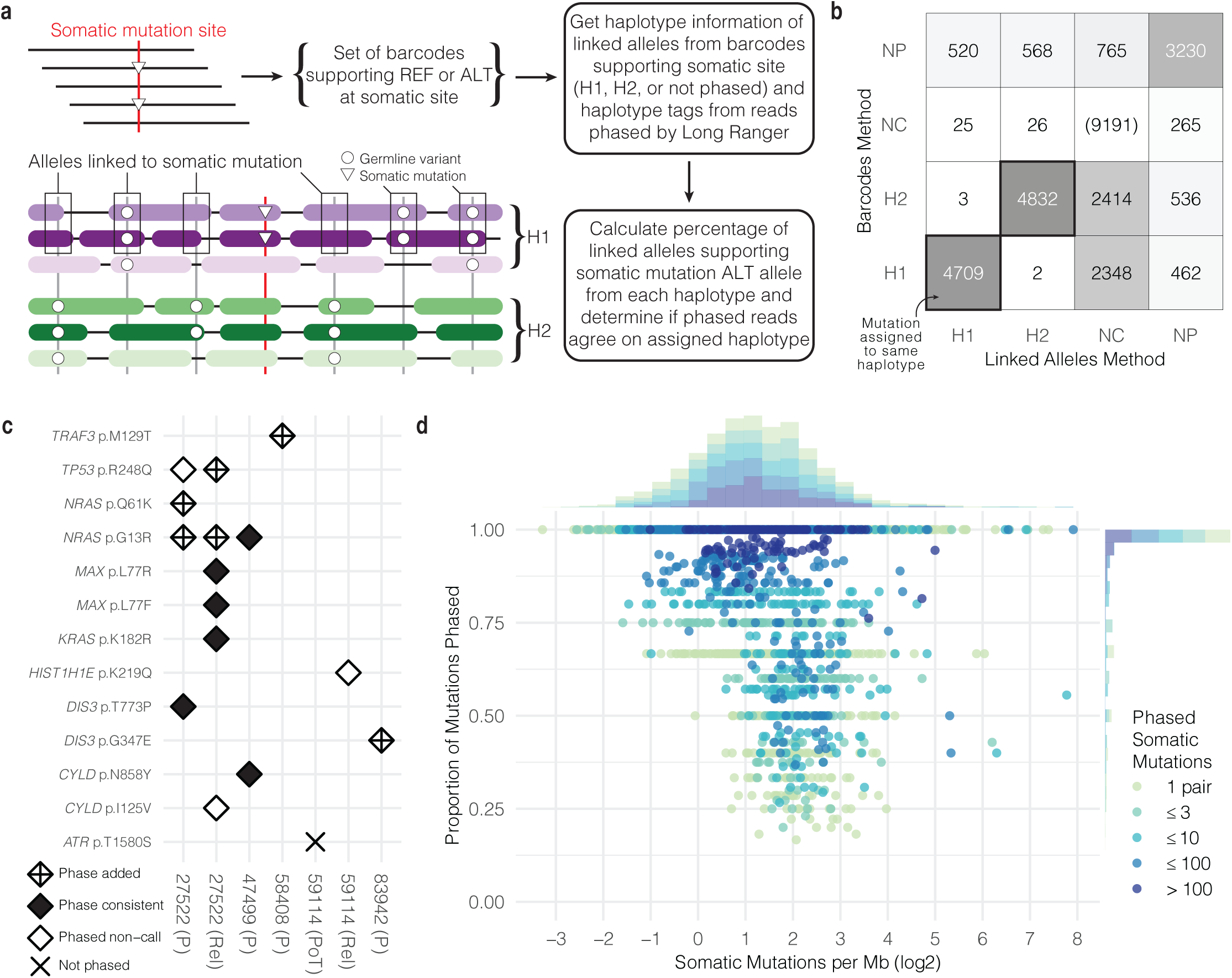
Phasing somatic mutations to haplotypes. **a.** Overview of methods used to phase somatic mutations. **b.** Number of somatic mutations phased using two phasing methods (H1 = phased to haplotype 1; H2 = phased to haplotype 2; NC = not enough coverage for phasing; NP = not phased). **c.** Phasing somatic mutations commonly observed in multiple myeloma. **d.** Distribution of somatic mutations per phase block and the proportion of mutations phased.

DNA, and we know the haplotype assignment of most (∼99%) linked alleles. We developed two methods to phase somatic mutations even if the mutation was not phased by the standard pipeline. In the “linked alleles” approach, if the linked alleles co-occurring with the somatic mutation are consistently phased to the same haplotype, we can infer the haplotype of the somatic mutation since it is most likely the same as the linked alleles. Alternatively, we can also use the “barcodes” approach which leans on the assigned phased of reads supporting the alternate allele as evidence. We required complete agreement of reads with assigned phases to confidently infer the haplotype of somatic mutations. In this approach, we extract the haplotype annotation for each read, which is reported as a tag in the phased bam output from Long Ranger. However, this information is not given for all reads. In our tumor sample data, 71.6% of reads overlapping a somatic mutation site were assigned a haplotype. Combining these two approaches increases phasing power when one approach lacks adequate coverage.

For six lrWGS samples with matched CD138+ sorted WGS, we called high-confidence somatic mutations using the sorted WGS tumor sample (see **Methods**). In total, we detected 32,842 somatic SNVs from our six sorted WGS samples, or 5,474 somatic SNVs per sample. Of those, 29,896 mutations (4,983 per sample) were SNVs with coverage in the matched lrWGS samples, and 20,705 (69.2%) met our minimum coverage requirement of at least 10 linked alleles from barcodes supporting the mutant allele or at least one phased read supporting the mutant allele. To establish a linked allele threshold at which we could confidently phase somatic mutations, we overlapped high-confidence somatic mutations from our WGS calls with phased Long Ranger calls to create a comparison set. Using the phased Long Ranger calls as the gold standard, we found that requiring at least 91% of linked alleles to be from the same haplotype before phasing a mutation led to an optimal balance of precision (0.997) and recall (0.936) (Supplementary Figure 4a) (see **Methods**). Overall, 79.4% (16,440/20,705 mutations) of somatic mutations with enough coverage were phased using that cutoff. Overall, the linked alleles and barcodes phasing methods were concordant on 99.95% of phasing decisions where both methods made a phasing decision (H1 or H2) (9,541/9,546 calls) (Fig. 2b). The barcodes approach added 5,760 calls where linked alleles did not have enough coverage or did not meet the phasing threshold. The linked alleles approach added 1,139 calls. See Supplementary Figure 4b for an overview of all results by phasing method.

We sought to contextualize the phasing performance of our simple heuristics focused on known somatic mutations within the broader landscape of genome-wide variant phasing software tools. We intersected variant phasing results reported by three tools (Long Ranger (v2.2.2), WhatsHap^49^ (v1.1), and HapCUT2^11^ (v1.3)) (see **Methods**) with our results to compare when each tool made a confident phasing decision. Of 20,705 variants with enough coverage, 34.0% (7,033/20,705 variants) were reported by each tool and were either phased or not phased. Our targeted, heuristic approach limited to known somatic mutations phased 88.2% (6,203/7,033 variants) in that intersection, while WhatsHap phased 59.3% (4,171/7,033 variants), HapCUT2 phased 52.0% (3,656/7,033 variants), and Long Ranger phased 52.0% (3,654/7,033 variants).

Figure 2c highlights seven samples with somatic mutations commonly associated with multiple myeloma, including mutations in *CYLD*, *DIS3*, *HIST1H1E*, *KRAS*, *NRAS*, and *TP53*.^45^ In 9 out of 16 examples shown, we confidently phased somatic mutations that were either not called or were not phased by Long Ranger. One mutation in *ATR* was not called by Long Ranger and was not phased by our approach since the linked alleles did not clearly favor one haplotype over the other (60.2% of phased linked alleles supporting the somatic mutation were phased to Haplotype 1, and 39.8% were phased to Haplotype 2). In 27522 (P), the *NRAS* G13R mutation was phased by our method to Haplotype 2, but was phased to Haplotype 1 in 27522 (Rel). However, since haplotype numbering is arbitrary, such differences are trivial. Further, we noticed that well-known hotspot *NRAS* mutations G13R and Q61K were both phased to the same haplotype in 27522 (P). Later analysis suggested that these two events occurred independently in separate tumor subclones.

We grouped high-confidence somatic mutations by phase block and found the proportion phased by our approach (Fig. 2d). The number of phased somatic mutations per megabase within each phase block showed a log2-normal distribution ranging from 0.10 to 241.3, with a median of 2.25. One application of phasing somatic mutations is establishing the pairwise haplotype relationship with other somatic mutations. Close to half of phase blocks longer than 1 kb had zero pairs of somatic mutations (44.8%, 2,212/4,941 phase blocks), with 11.1% having zero somatic mutations and 33.6% having only one somatic mutation. But among those 2,729 phase blocks longer than 1 kb with at least one pair of somatic mutations, 33.2% had exactly one pair, 18.0% 2-3 pairs, 20.4% 4-10 pairs, 22.3% 11-100 pairs, and the remaining 6.0% had more than 100 pairs. 64.6% of those phase blocks had every mutation phased, and 77.5% had at least 75% of mutations phased.

### Pairs of phased somatic mutations illustrate patterns of clonal evolution

In short read sequencing, if two mutant alleles are called together on the same read or read pair, then we can infer they occurred in the same cell and on the same molecule of DNA. With lrWGS, we have the benefit of more linked-reads to consider when we look for such co-occurring mutations. From six samples with lrWGS as well as high-confidence somatic mutation calls and CNV profiles from WGS, we focused on mutations in copy number neutral regions with coverage between 10 and 100 phased barcodes at that position and one or more barcodes supporting the alternate allele (Supplementary Figure 5a). We examined 59,063 pairs of mutations and, as expected, the probability of one barcode covering both sites decreases as the distance between sites increases, with 98.4% (54,643/55,559 pairs) of mutation pairs located greater than 62 kb apart sharing no overlap. (62 kb is the median of the mean molecule lengths described in Fig. 1d.) Therefore, we focused on the 3,504 mutation pairs within 62 kb. For the 2,648 mutation pairs within this proximity but greater than 100 bp apart, 13.0% did not share any barcodes, 77.3% shared between 1 and 10 barcodes, 8.3% between 11 and 20 barcodes, and 1.4% greater than 20 barcodes (Fig. 3a). For the 856 mutation pairs located less than 100 bp apart, each pair had at least one shared barcode (Supplementary Figure 5b). Overall, 5.9% (3,504/59,063 pairs) of somatic mutation pairs were within 62 Kb (Fig. 3b). Of those, 90.2% (3,159/3,504 pairs) share at least one barcode in common, and, of those, 64.6% (2,042/3,159 pairs) have a barcode on which one or both somatic mutations is represented, potentially enabling direct observation of mutation patterns in the same cell.

**Figure 3.**
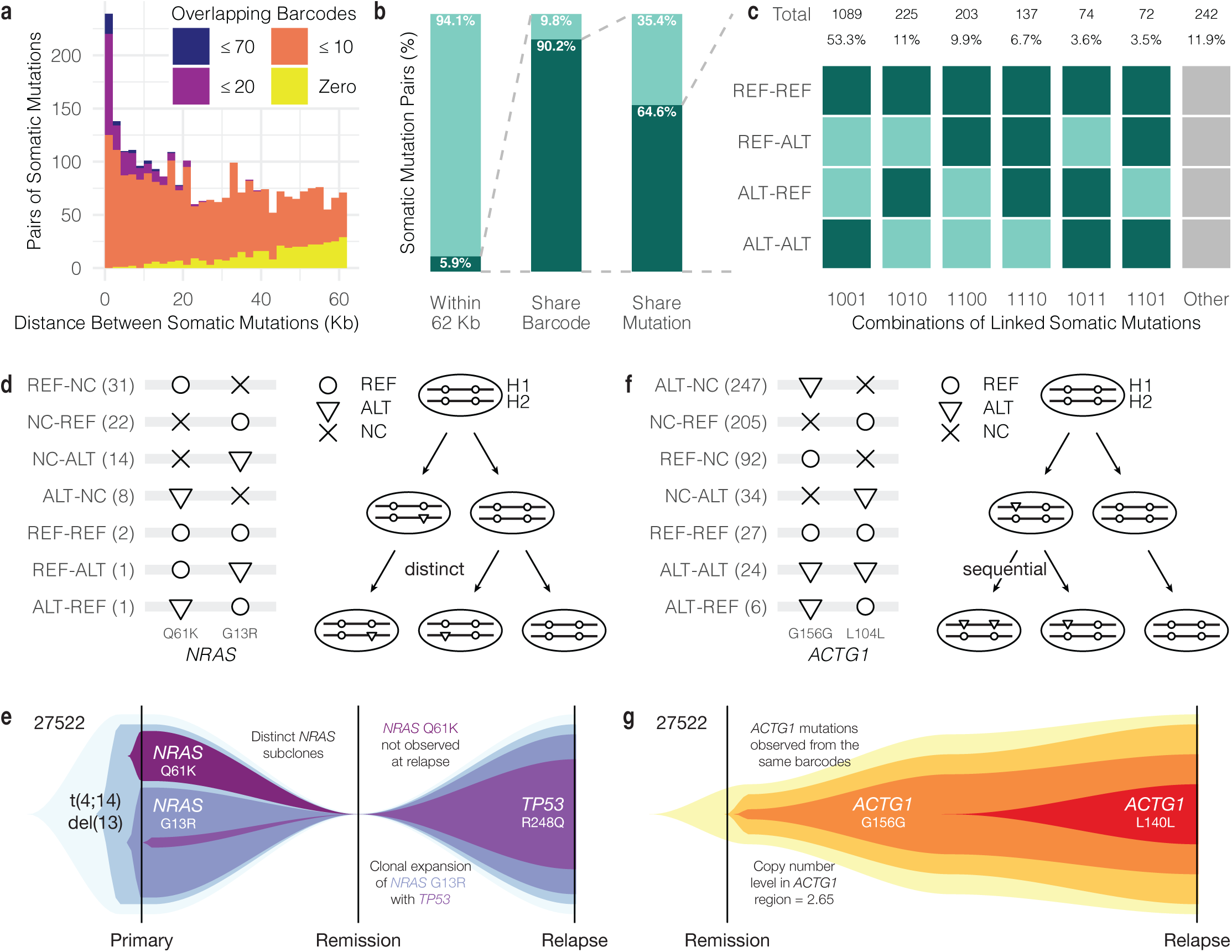
Tumor evolution models derived from mutation pairs. **a.** Number of overlapping barcodes by distance between somatic mutations. **b.** Proportion of somatic mutation pairs in close proximity sharing barcodes and mutations. **c.** Patterns of mutation pairs observed on barcodes (REF = reference allele; ALT = alternate allele). A dark green square indicates that a barcode with that pattern of two alleles was observed. Combinations of patterns can interpreted as evidence of sequential (e.g. 1101, 1011) or distinct (e.g. 1110) mutations. **d.** *NRAS* mutation pair observed in 27522 (P) and evolution model (NC = no coverage). **e.** Interpretation of evolution model observed from *NRAS* mutation pair in 27522 (P). **f.** *ACTG1* mutation pair observed in 27522 (Rel) and evolution model. **g.** Interpretation of evolution model observed from *ACTG1* mutation pair in 27522 (Rel).

We then considered the observed pairwise relationship of each reference and alternate allele on barcodes covering the two somatic sites (Fig. 3c). Of the 2,042 remaining mutation pairs, most (53.3%) share barcodes that only support either both reference alleles (REF-REF) or both alternate alleles (ALT-ALT). This means they have at least one barcode where both alleles are REF and at least one barcode where both alleles are ALT. Other observed patterns are less common, but include REF-REF with REF-ALT or ALT-REF, in which there is at least one barcode supporting one of the alternate alleles but not both. 6.7% of pairs show barcodes supporting REF-ALT and ALT-REF. In these cases, if the two alternate alleles are phased to the same haplotype in a copy number neutral context, this could indicate that the two mutations occurred on the same haplotype but in different cells. Finally, 7.1% of pairs have a pattern of REF-ALT or ALT-REF along with ALT-ALT, suggesting a pattern of sequential mutation events. With greater tumor purity, we would expect to see a higher proportion of informative allele patterns with the potential to inform patient-specific models of tumor evolution.

One such example where the pattern of somatic mutations may be informative for refining tumor phylogenies and may have clinical implications came from CD138+ sorted sample 27522 (P). We observed two hotspot mutations in *NRAS* (G13R and Q61K) (Fig. 3d). *NRAS* is a known cancer driver oncogene and mutations may lead to dysregulation of the Ras pathway. We phased both mutations to the same haplotype (H2) (Supplementary Figure 6). We observed 2 barcodes supporting REF-REF, 1 barcode supporting REF-ALT, and 1 barcode supporting ALT-REF. Based on sorted lrWGS data, the variant allele frequency (VAF) of the G13R mutation was 35.7% and the Q61K VAF was 22.2% at the primary stage. At relapse, the G13R VAF was 20.5% and the Q61K mutation was not detected (VAF 0.0%). Such basic VAF calculations must be interpreted within the context of imperfect tumor cell sorting, tumor heterogeneity with subclonal structure, and potential partial copy number loss on the opposite haplotype (Supplementary Figure 3, Supplementary Figure 6). It may be clinically relevant to know if the two mutations occurred independently or in the same subclone even though multiple activating mutations in the same gene are not necessary for clonal expansion. Without the benefit of phasing, one possible interpretation could be that Q61K occurred in the same clone as G13R, and then the double mutant subclone was eliminated after therapy. However, with linked-reads, we directly observed both mutations occurring without the other, and we never observed them together, guiding the interpretation that these mutations occurred independently in separate subclones and that the Q61K subclone was later lost (Fig. 3e).^50^

In another instance, we detected a pair of mutations in *ACTG1* (G156 and L104) that may have occurred in sequential order on the same biological haplotype. Six barcodes demonstrate the ALT-REF pattern, with ALT G156 and REF L104, and 24 barcodes had ALT-ALT with both sites mutated (Fig. 3f). Under a parsimonious model in which the same mutation occurs only once, the G156 mutation must have preceded the L104 mutation. Since there are barcodes supporting both mutant alleles simultaneously, the mutations most likely occur within the same cells, and we interpret this to mean the cells with both mutations form a later subclone within the subclone of cells with only the G156 mutation (Fig. 3g). We also noted elevated copy number in this region (estimated to be 2.65). This would often preclude clonality analysis due its effect on the VAF.^51^ However, the combination of alleles present on the same barcodes enables us to interpret a sequential order of events.

### Oncogenic IGH translocations in myeloma map to specific haplotypes

Multiple myeloma is characterized by recurrent clonal translocations that take advantage of overexpressed IGH locus by dysregulating oncogene expression. Barwick, et al. analyzed 795 newly-diagnosed multiple myeloma patients from the Multiple Myeloma Research Foundation CoMMpass study (NCT01454297) and reported clonal translocations across the cohort, including 16% of patients with t(11;14) impacting *CCND1*, 11% with t(4;14) (*WHSC1*), 3.3% with t(14;16) (*MAF*), 1.1% with t(6;14) (*CCND3*), and 1.0 % with t(14;20) (*MAFB*).^52^ In our cohort of 14 patients, we detected common myeloma translocations from lrWGS using the Long Ranger pipeline and as well as from sorted WGS in 9 matching samples and found t(11;14) in 2 patients and t(4;14) in 1 patient (see **Methods**).^53^ After selecting high-confidence events reported from sorted WGS, we found supporting evidence from lrWGS barcodes and mapped those events to haplotypes.

From the 9 matched sorted WGS samples, we identified 88 high-confidence translocations (see **Methods**). We then interrogated matching lrWGS data to find barcodes supporting the event. Of those 88 high-confidence events, 20.5% (18/88 events) had at least two barcodes with a read pattern in support of the translocation. This low rate of support may be attributed to most lrWGS samples not being sorted to select for tumor cells. However, of the 18 events with at least two barcodes, the read haplotype assignment of 94.4% (17/18 events) of translocations showed a consistent haplotype assignment, suggested that using high-confidence SV calls from WGS is a robust prior for haplotype mapping of SVs in high purity lrWGS data.

In Patient 27522, 6 out of 7 SVs detected from both Primary and Relapse samples were also detected from WGS (Supplementary Figure 7a). This patient had a t(4;14) event detected at primary diagnosis present later at relapse which juxtaposed the IGH enhancers with *WHSC1* and *FGFR3*, leading to overexpression of both oncogenes (Supplementary Figures 7b, 8a-b). *WHSC1* overexpression in t(4;14) tumors increases dimethylation of H3K36 and broadly dysregulates the myeloma epigenome.^54^ The coverage heat map showing where discordant barcodes map on chr4 and chr14 clearly shows the translocation breakpoint within the first intron of *WHSC1* at chr4:1871962 and near *IGHM* on chr14 and also indicates a deletion proximal to the translocation breakpoint on chr14. We then visualized the coverage pattern of barcodes with reads mapping to both chromosomes in a window around the reported t(4;14) breakpoints (Supplementary Figure 7c). The barcode coverage indicates a reciprocal event leading to two new derived chromosomes der(4) and der(14) with reads from barcodes supporting t(4;14) arbitrarily assigned to H2 on both chromosomes. A pair of events in 27522, t(6;17) and t(4;6), showed similar breakpoints on chr6, approximately 14 kb apart. However, we did not observe convincing evidence of barcodes with a read coverage pattern linking the three chromosomes, supporting the interpretation that these events occurred independently.

For 77570 (P), Long Ranger reported two t(11;14) events affecting different regions of IGH but with the same breakpoint upstream of *CCND1* (Supplementary Figure 7d-e, 8c-f). One event linked the IGH variable gene region (chr14:106269142) to *CCND1* on chr11. The other at chr14:105741942 linked the coding region of *IGHG1* to the same *CCND1* breakpoint. Barcode coverage analysis suggests these two reported events may actually be one complex reciprocal event with a t(11;14) translocation and deletion on chr14 giving the observed pattern of read coverage upstream and downstream of each breakpoint (Supplementary Figure 7f).

One application of translocation mapping is matching allele-specific expression to translocation events, for example if a germline heterozygous coding variant from the same haplotype of the dysregulating translocation were detected from RNA-seq, then the connection between translocation and expression could be made more explicitly.

### Shared germline variants from matched samples enable phase block extension

Phase block boundaries may differ between samples originating from the same patient. However, samples from the same patient do share germline variants, and those germline variants should be phased together in the same groups in both samples.^55^ In contrast to previous sections in which somatic mutations from the same sample and same phase block were analyzed together, by comparing the phase of germline variants from overlapping phase blocks from two samples, we can determine if the two phase blocks are oriented the same way, or if one needs to be flipped for them to be consistent. We compared germline variants from overlapping phase blocks found in two samples, the target sample and the reference sample (Fig. 4a) (see **Methods**). If the shared germline variants were consistently assigned to the same haplotype, the target and reference phase blocks have the same orientation. If they were consistently assigned to opposite haplotypes, they have opposite orientation and the target needs to be switched. If two target phase blocks overlap the same reference phase block, then we can infer the haplotype orientation of the target phase blocks. However, if a switch error occurs in one phase block, that error will propagate as phase blocks are extended.

**Figure 4.**
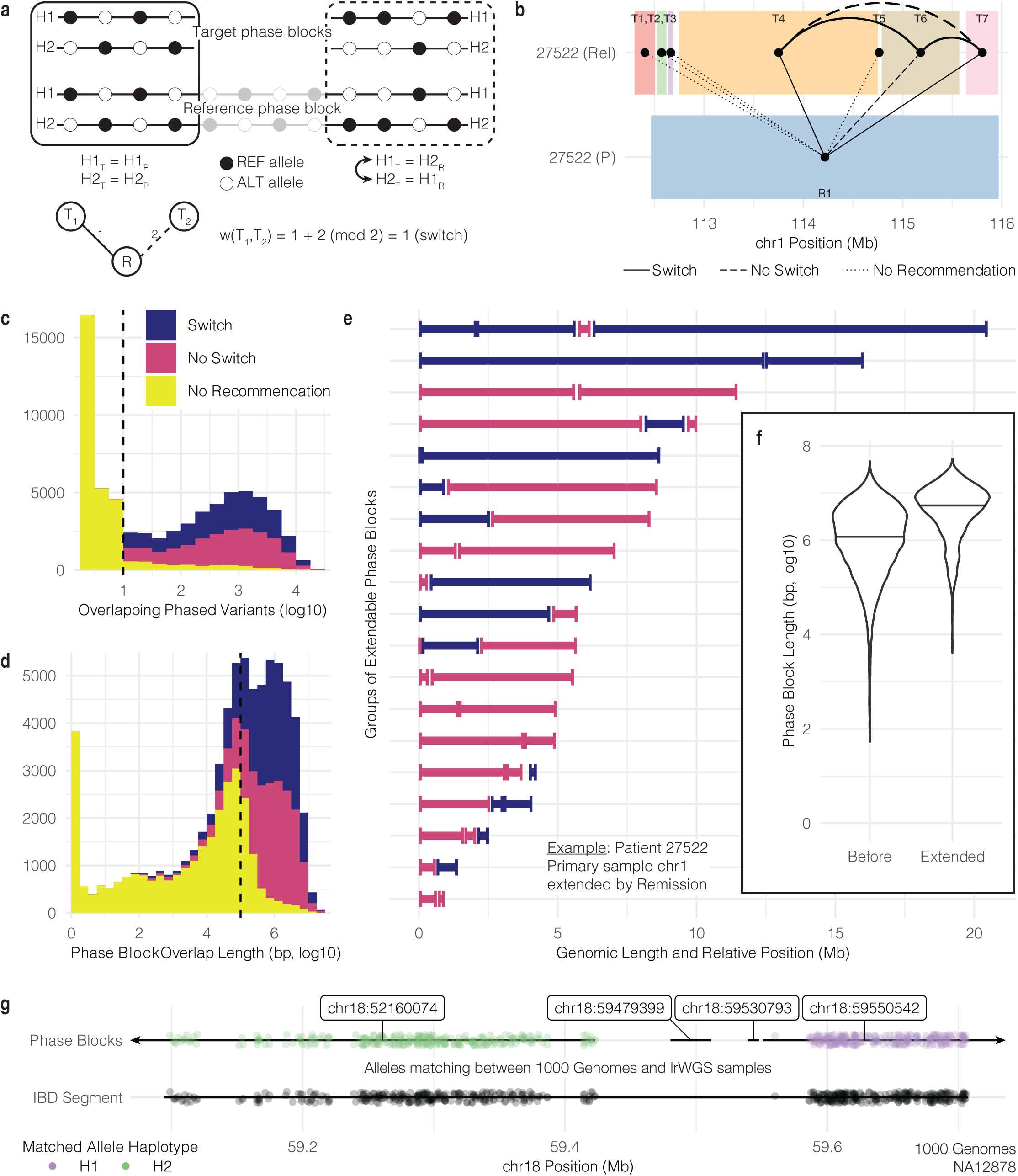
Extension of phase blocks using additional sample information. **a.** Model for phase block extension using overlap between target and reference phase blocks. **b.** Data-driven example of phase block overlap between samples. **c.** Number of phased variants needed for switch/no switch recommendation. **d.** Length of phase block overlap needed for switch/no switch recommendation. **e.** Phase block groups extended by overlap with another sample. **f.** Distribution of phase block lengths before and after extension. Violin plots defined as: center line, median; violin limits, minimum and maximum values; individual points not shown. **g.** Use of identity-by-descent segments as overlap between phase blocks.

We analyzed data from 6 patients having multiple tumor samples, with a total of 68,374 overlapping phase blocks from 26 target and reference sample pairs. For example, we examined phase blocks originating from chr1 of 27522 (P) and 27522 (Rel), using 27522 (P) as the reference sample (bottom) and 27522 (Rel) as the target sample (top) (Fig. 4b). Reference phase block 1 (R1) (colored blue) spans multiple target phase blocks (T1-T7). For T1, T2, T3, and T5, there are not enough overlapping variants to draw conclusions about their orientation relative to R1. Phase blocks T4 and T7 must be switched in order to be consistent with R1, and T6 is already consistent with R1. Since T4 and T7 have the same orientation relative to R1, they have the same haplotype orientation and do not need to be switched. However, T6 must be switched to be consistent with T4 and T7. By grouping disconnected phase blocks together, we increase the number of pairs of loci with known haplotype orientation.

In general, at least 10 overlapping phased variants are required before making a switch or no switch recommendation (Fig. 4c). Since the number of shared variants is correlated with the length of the overlap, the length of overlap tends to be greater than 100 kb before a recommendation can be made (Fig. 4d). We were not surprised to find roughly equal proportions of recommendations to switch (28.3%) and not switch (27.6%) given that haplotype numbering is random. For the remaining 44.1% of cases, the algorithm was not able to make a recommendation to switch or not switch. For extendable phase blocks from chr1 in target sample 27522 (P) (extended by reference 27522 (Rem)), we found that, before extension, the median phase block length was 1.6 Mb, and after extension, it was 5.7 Mb, a 3.5-fold increase (Fig. 4e). Similarly, from all samples with extendable phase blocks, we found that median phase block length increased from 1.2 Mb (6.1 on log10 bp scale) to 5.5 Mb (6.7 on log10 bp scale), a 4.6 fold increase from before extension to after extension (Fig. 4f).

We also developed methods to leverage publicly available population-scale phased data to learn more about the origin of haplotypes present in our cohort and to improve our lrWGS results. After reporting identical-by-descent (IBD) segments shared between 2,504 individuals from 1000 Genomes data (see **Methods**; see **Data availability**), we identified IBD segments overlapping multiple lrWGS phase blocks in NA12878.^56^ Using phased heterozygous variants shared between the 1000 Genomes VCF of this sample and the VCF output from Long Ranger, we found the proportion of IBD alleles matching each haplotype in each phase block. IBD alleles consistently matched one haplotype or the other with the occasional short switch error. For example, NA12878 shares an IBD segment with NA10851 from position 59,094,547 to 59,706,930 on chr18 (LOD score 15.64, 1.576 cM) (Fig. 4g). That IBD segment bridges multiple lrWGS phase blocks. Since the IBD alleles match Haplotype 2 from phase block chr18:52160074 and match Haplotype 1 from chr18:595505042, those two phase blocks may be in opposite orientation.

## Discussion

As sequencing technologies evolve and analysis methods more regularly include haplotype phasing, somatic mutation phasing will become a more common practice. The current methodological approaches to haplotype-aware somatic mutation analysis will mature from *ad hoc* investigations to standard pipelines. We have developed a systematic approach to somatic mutation analysis in a cohort of multiple myeloma patients over the course of disease. Our methods build on the backbone of the Long Ranger variant calling and phasing pipeline for linked-read sequencing data. These methods are an opportunity for future development in a climate of rapid technological advances with many applications. We need better understanding of the haplotypes carrying germline variants related to predisposition of many diseases, including cancer, as well as better methods to identify ancestry-specific risk modifiers.^57–63^ Biallelic *TP53* inactivation indicates poor prognosis in multiple myeloma^64^, and double *PIK3CA* mutations on the same haplotype can be more oncogenic but also more susceptible to PI3Ka inhibitors.^65^ Other medical applications of linked-read sequencing include more sensitive prenatal diagnosis ^66,67^, better predictions about how protein structure may change in response to multiple mutations ^68^, and more accurate neoepitope prediction.^69^ Tools such as HAPDeNovo capitalize on haplotype structures from linked-reads to eliminate false-positive from studies of rare, *de novo* variation.^70^

We noted several limitations in our analysis potentially due to data generation. We observed shorter phase blocks in our skin normal controls samples potentially due to lower input molecule size or sequencing depth. For our somatic analyses, an important caveat was controlling for copy number changes which disrupt the strict two haplotype paradigm of variant phasing. Another limitation of our somatic analysis was low tumor purity. Only 4 of our 23 samples were CD138+ sorted, and two samples in particular gave us the most confident results. Higher tumor purity and lower variability in cell-type composition are likely important for robust somatic variant haplotype analysis. Calling somatic mutations with low variant allele frequency is a challenge for any mutation caller, especially those like Long Ranger built for germline variant detection. In our case, pairing linked-read data with high-confidence somatic mutation calls from a separate WGS sample was necessary to gain sensitivity. Future analyses using lrWGS in multiple myeloma should include analysis of chromoplexy and chromothripsis as these complex events are important in MM pathogenesis but cannot be fully appreciated using short reads.^71^ Additionally, long-range PCR of known somatic variant regions could validate the phasing performance and data interpretations enabled by our framework.

Moving beyond next-generation sequencing to Third Generation and single-cell approaches holds the promise of increased resolution in cancer genome analyses.^72–75^ With long reads and linked-reads, we get haplotype resolution. With single-cell RNA-seq, we observe cell-specific patterns of gene expression and copy number and can map coding mutations to specific cells.^76^ Single-cell DNA sequencing analyses, including approaches that incorporate haplotypes, offer even deeper resolution of tumor evolution and the ability to optimize treatment strategies.^18,77–84^ Methodological integration of single-cell data with the resolution gained from haplotype analysis is a direction for continued research.

## Methods

### SomaticHaplotype modules

Our framework of interconnected modules builds on the phased bam and variant output files from the Long Ranger pipeline (10X Genomics). Our analysis code was written in python and is freely available under the MIT license (see **Code availability**). Additional inputs to our pipeline may include high-confidence somatic mutation calls and identity-by-descent segments. There are five analysis modules: *phaseblock*, *summarize*, *somatic*, *extend*, and *ancestry*.

#### Phaseblock

The *phaseblock* module is the first module run on any new data. The inputs are a phased bam and phased variant call format (VCF) file from the Long Ranger pipeline. Given a genomic range of interest, such as an entire chromosome, *phaseblock* constructs PhaseBlock and Variant objects by extracting information from reads and variant calls. PhaseBlock objects collect information about variants common to phase blocks identified by Long Ranger. Variant objects store information about small variants, including genotype and phase, and map to specific PhaseBlock objects based on tags given by Long Ranger. Each Variant also stores the barcodes of reads supporting the reference and alternate allele at that position. The objects are designed with methods for later utility. Dictionaries referencing those objects are stored in an output file used as input to downstream modules.

#### Summarize

The *summarize* module takes input from *phaseblock* and produces a summary of each phase block and a global summary about phase block lengths. Output from *summarize* is used as input to *somatic* and *extend*.

#### Somatic

The *somatic* module collects barcode and haplotype information supporting somatic mutation sites. Somatic mutation sites are defined using an input parameter, either as a mutation annotation format (MAF) file or list of mutations. Barcodes supporting somatic mutation sites are extracted by a separately-run submodule called 10Xmapping which mines the bam for reads supporting the mutation site and also from the VCF if the mutation was called by Long Ranger. The 10Xmapping submodule identifies bam reads supporting the reference and alternate alleles at a somatic mutation site and gathers barcode and haplotype information from each read. 10Xmapping is contained as a submodule of in our repository but is also freely available at https://github.com/ding-lab/10Xmapping. Output from *somatic* includes information about every (germline and somatic) variant from barcodes overlapping each somatic mutation site, information necessary for phasing each somatic mutation, barcode sharing analysis of each pair of somatic mutations, and somatic mutation summaries for each phase block.

In later analysis, users interpret output from *somatic* to decide if somatic mutations are phased or not. For example, we combined two approaches to determine the phase of each somatic mutation. In our “linked alleles” approach, we analyzed the proportion of linked alleles mapping to a particular haplotype and found 0.91 (and above) to be an appropriate threshold that balanced phasing decision precision and recall. We combined that with the “barcodes” approach, which relies on the reported haplotype assignment of reads supporting the somatic mutation. We determined a somatic mutation to be phased if at least one barcode supported the mutant allele and all barcodes supporting the mutant allele agreed on the haplotype assignment. For pairs of somatic mutations, the barcode sharing analysis finds barcodes with reads mapping to both somatic mutation sites. For each barcode, the alleles supporting each site are combined as allele pairs (REF-REF, REF-ALT, ALT-REF, and ALT-ALT).

#### Extend

The extend module combines germline variants from two related samples (e.g. from the same individual) to determine the haplotype orientation between disconnected phase blocks in one of the samples. Once the haplotype orientation between two phase blocks is determined, the phase blocks can be conceptually extended. The two samples are defined as the “target” (with phase blocks to be extended) and the “reference”, which the target is compared against. To determine if the target and reference phase blocks have the same or different haplotype orientation, *extend* compares the haplotype assignments of overlapping germline variants and finds the proportion of target haplotype assignments that need to be switched in order to be consistent with the reference. *Extend* uses a two-sided binomial test (significant number of “switch” or “not switch” given a conservative switch error rate) and a hard cutoff (more than 95% “switch” or less than 5% “switch”) to determine if the target and reference phase blocks have the same or opposite orientation. Then *extend* module then builds a bipartite graph in which nodes are phase blocks and edges connect overlapping target and reference sample phase blocks. Edge weights are defined as 1 if a switch is necessary between the target and reference phase block or 2 if a switch is not necessary. If two target phase blocks overlap the same reference phase block, then there is a connected path between the target phase blocks and we find the sum of the weighted edges connecting them. If the sum (mod 2) is zero, then the two target phase blocks have the same orientation. If the sum (mod 2) is one, then they have opposite orientation. *Extend* output describes the overlap of each target phase block with reference phase blocks and also forms groups of connected target phase blocks that may be extended via this method.

#### Ancestry

The *ancestry* module uses a similar concept to *extend* but instead relies on output from an identity-by-descent tool such as Refined IBD instead of phase blocks from a related sample.^56^ By examining the haplotype assignment of alleles from overlapping IBD segments and phase blocks defined by Long Ranger, *ancestry* may bridge gaps between phase blocks and find where phase block haplotype orientations are congruent or not. *Ancestry* also assigns population history to portions of phase blocks that overlap IBD segments.

### Generation of linked-read whole genome sequencing data

The 10X Genomics Chromium System generates linked-read sequencing data. From a bulk sample of cells, long fragments of DNA, also called high-molecular weight (HMW) DNA, are isolated into an individual gel bead in emulsion (GEM). Each GEM contains a gel bead with primers including a 16-bp DNA barcode unique to that GEM. The gel bead dissolves and releases the barcoded primers, which attach to the DNA and undergo isothermal amplification. Now each short fragment of amplified DNA contains a barcode identifying which GEM it originated from. The GEMs break and the barcoded fragments are pooled together and sequenced.

### Patient cohort

Fourteen (10 male, 4 female) patients with multiple myeloma were included in the analysis. The median age at diagnosis was 63 (range 46-69). Eight patients had IgG isotype (4 kappa and 4 lambda), 2 had IgA kappa isotype, 2 had light chain only disease (1 kappa and 1 lambda), and 2 were non-secretory. Five were International Staging System Stage I, 2 were Stage II, 3 were stage III, and 4 were unreported. The median plasma cell burden by flow cytometry in bone marrow at diagnosis was 24% (range 4-63). By standard fluorescence in situ hybridization (FISH), 1 patient had t(4;14), 3 had t(11;14), and 2 showed del(17p). A total of 23 samples were collected from multiple disease stages, including smoldering multiple myeloma (SMM), primary diagnosis, pre- and post-transplant, remission, and relapse.

### Sample collection and data generation

Research bone marrow aspirate samples were collected at the time of the diagnostic procedure. Bone marrow mononuclear cells (BMMCs) were isolated using Ficoll-Paque. BMMCs were cryopreserved in a 1:10 mixture of dimethyl sulfoxide and fetal bovine serum. Upon thawing, whole BMMCs were used for linked-read whole genome sequencing. Plasma cells were separated from a sub-aliquot by positive selection using CD138-coated magnetic beads in an autoMACs system (Miltenyi Biotec, CA) and used for whole genome and exome sequencing. Skin punch biopsies were performed at the time of the diagnostic bone marrow collection to serve as normal controls. Although many studies use peripheral blood mononuclear cells (PBMCs) as a control, abnormal B cells and circulating tumor cells frequently contaminate the peripheral blood of patients with multiple myeloma. Therefore, using PBMCs may lead to the omission of genetic events potentially important in disease pathogenesis.

#### Linked-read whole genome sequencing (lrWGS)

Normal skin samples were processed with a standard Qiagen DNA isolation kit resulting in 10-50Kb DNA fragments. 250K tumor cells were processed with the MagAttract HMW DNA extraction kit (Qiagen) resulting in 100-150Kb DNA fragments. 600-800ng of normal DNA was size selected on the Blue Pippin utilizing the 0.75% Agarose Dye-Free Cassette to attempt to remove low molecular weight DNA fragments. The size selection parameters were set to capture 30-80 Kb DNA fragments (Sage Science). The resulting size selected DNA from the normal samples and the HMW DNA from the tumor cells were diluted to 1ng/μL prior to the v2 Chromium Genome Library prep (10X Genomics).

Approximately 10-15 DNA molecules were encapsulated into nanoliter droplets. DNA molecules within each droplet were tagged with a 16 nucleotide barcode and 6 nucleotide unique molecular identifier during isothermal incubation. The resulting barcoded fragments were converted into a sequence ready Illumina library with an average insert size of 500bp. The concentration of each library was accurately determined through qPCR (Kapa Biosystems) to produce cluster counts appropriate for sequencing on the HiSeqX/NovaSeq6000 platform (Illumina). 2×150 sequence data were generated targeting 30x (normal) and 60x (tumor) coverage providing linked-reads across the length of individual DNA molecules.

#### Standard whole genome sequencing (WGS)

Manual libraries were constructed with 50-2000ng of genomic DNA utilizing the Lotus Library Prep Kit (IDT Technologies) targeting 350bp inserts. Strand-specific molecular indexing is a feature associated with this library method. The molecular indexes are fixed sequences that make up the first 8 bases of read 1 and read 2 insert reads. The concentration of each library was accurately determined through qPCR (Kapa Biosystems). 2×150 paired-end sequence data generated ∼200 Gb per tumor sample leading to 60x (tumor) haploid coverage.

### lrWGS data processing with Long Ranger

Long Ranger (10X Genomic) performs linked-read alignment, variant calling, and variant phasing. We ran Long Ranger (v2.2.2) to align reads to the human genome reference GRCh38 (GRCh38-2.1.0) and used --vcmode with GATK^85,86^ (version 3.7.0-gcfedb67) for variant calling. Long Ranger also produces quality metrics associated with each sample. Publicly-available 1000 Genomes lrWGS samples were processed with Long Ranger (version 2.2.1) and aligned to hg19.

### lrWGS data processing with other tools

In addition to Long Ranger, we used WhatsHap^49^ (v1.1) and HapCUT2^11^ (v1.3) to phase our linked read WGS samples using human genome reference GRCh38 (GRCh38-2.1.0). We applied the additional extractHAIRS and LinkFragments steps to prepare our 10X data for use by HapCUT2.

### High-confidence somatic mutation detection

Somatic mutations were called by our SomaticWrapper pipeline, which includes four established bioinformatic tools, namely Strelka^87^, Mutect^88^, VarScan2^89^ (2.3.83), and Pindel^90^ (0.2.54). We retained SNVs and INDELs using the following strategy: keep SNVs called by any 2 callers among Mutect, VarScan, and Strelka and INDELs called by any 2 callers among VarScan, Strelka, and Pindel. For these merged SNVs and INDELs, we applied coverage cut-offs of 14X and 8X for tumor and normal, respectively. We also filtered SNVs and INDELs with a high-pass variant allele fraction (VAF) of 0.05 in tumor and a low-pass VAF of 0.02 in normal. The SomaticWrapper pipeline is freely available at https://github.com/ding-lab/somaticwrapper.

### Copy number profiling

We used BIC-seq2^91^, a read-depth-based CNV calling algorithm to detect somatic copy number variations (CNVs) using standard WGS tumor samples and paired skin linked-read WGS data. The procedure involves 1) retrieving all uniquely mapped reads from the tumor and paired skin BAM files, 2) removing biases by normalization (NBICseq-norm_v0.2.4) 3) detecting CNV based on normalized data (NBICseq-seg_v0.7.2) with BIC-seq2 parameters set as -- lambda=90 --detail --noscale --control. We defined copy number neutral regions as having a log_2_ copy number ratio between -0.25 and 0.2 in the sorted WGS.

### Tumor purity estimation

We used the R package sciClone^51^ (v1.1.0) to estimate tumor purity based on clusters detected using variants from copy number neutral regions. We designated the cluster with the greatest median variant allele frequency (VAF) as the founding clone and doubled its VAF to estimate the sample’s tumor purity.

### Structural variant detection

Somatic structural variants (SVs) were detected by Manta^53^ using tumor/normal sample pairs of standard WGS and paired skin linked-read WGS. SVs were filtered according to the following guidelines. Record-level filters included a QUAL score < 20; somatic variant quality score < 30; depth greater than 3x the median chromosome depth near one or both variant breakpoints; for variants significantly larger than the paired read fragment size, no paired reads support the alternate allele in any sample. Sample-level filters included a Genotype Quality < 15. This approach optimizes the analysis of somatic variation in tumor/normal sample pairs. In addition to the built-in Manta filters (labeled as PASS), we further prioritized the high-confidence variants by (1) the number of support spanning read pairs >= 5; (2) the coverage at the given breakpoints > 10; (3) events must involve only autosomes and/or sex chromosomes; (4) events passing manual IGV review on the read evidence. We also used gemtools^33^ (https://github.com/sgreer77/gemtools) and the python package pysam (0.15.3) with samtools^92^ (v1.9) to identify reads and barcodes supporting SVs in lrWGS.

### Identity-by-descent reporting

We obtained phased haplotype information for 2,504 individual from the 1000 Genomes and ran Refined IBD with default parameters (refined-ibd.16May19.ad5) (see **Data availability**).^56^

## Data availability

The Washington University Institutional Review Board approved the study protocol, and all relevant ethical regulations, including obtaining informed consent from all participants, were followed. Patients were treated and sampled at Washington University in St. Louis.

All data and scripts necessary to recreate figures are available at doi.org/10.6084/m9.figshare.12295922.

Publicly-available 1000 Genomes lrWGS samples can be downloaded from https://support.10xgenomics.com/genome-exome/datasets/2.2.1/NA12878_WGS_v2 and https://support.10xgenomics.com/genome-exome/datasets/2.2.1/NA19240_WGS_v2.

Phased 1000 Genomes VCFs (2,504 samples) were downloaded from http://ftp.1000genomes.ebi.ac.uk/vol1/ftp/release/20130502/.

The remaining data and methods are available in the Article, Supplementary Tables, or are available from the author upon reasonable request.

## Code availability

SomaticHaplotype is freely-available at https://github.com/ding-lab/SomaticHaplotype.

## Acknowledgements

This work has been supported by the Paula C. and Rodger O. Riney Blood Cancer Research Initiative Fund to L.D. and R.V. and NCI U24CA211006 and U2CCA233303 funds to L.D.

## Author contributions

L.D. and R.V. led project design. S.M.F. led tool development, performed data analysis, wrote manuscript, generated figures. N.V.T., Y.L., Q.G., L.Y., and H.S. ran tools for alignment, mutation, SV, and CNV calling. A.W., Q.G., G.D., M.S., S.C., and R.G.J. contributed to tool development. R.S.F. and C.C.F. led sequencing data generation. J.K., D.R.K., and M.A.F. managed in-house sample collection. R.G.J., K.C., J.F.D., R.V., and L.D. reviewed the manuscript.

## Supplementary Figure Legends

**Supplementary Figure 1.**
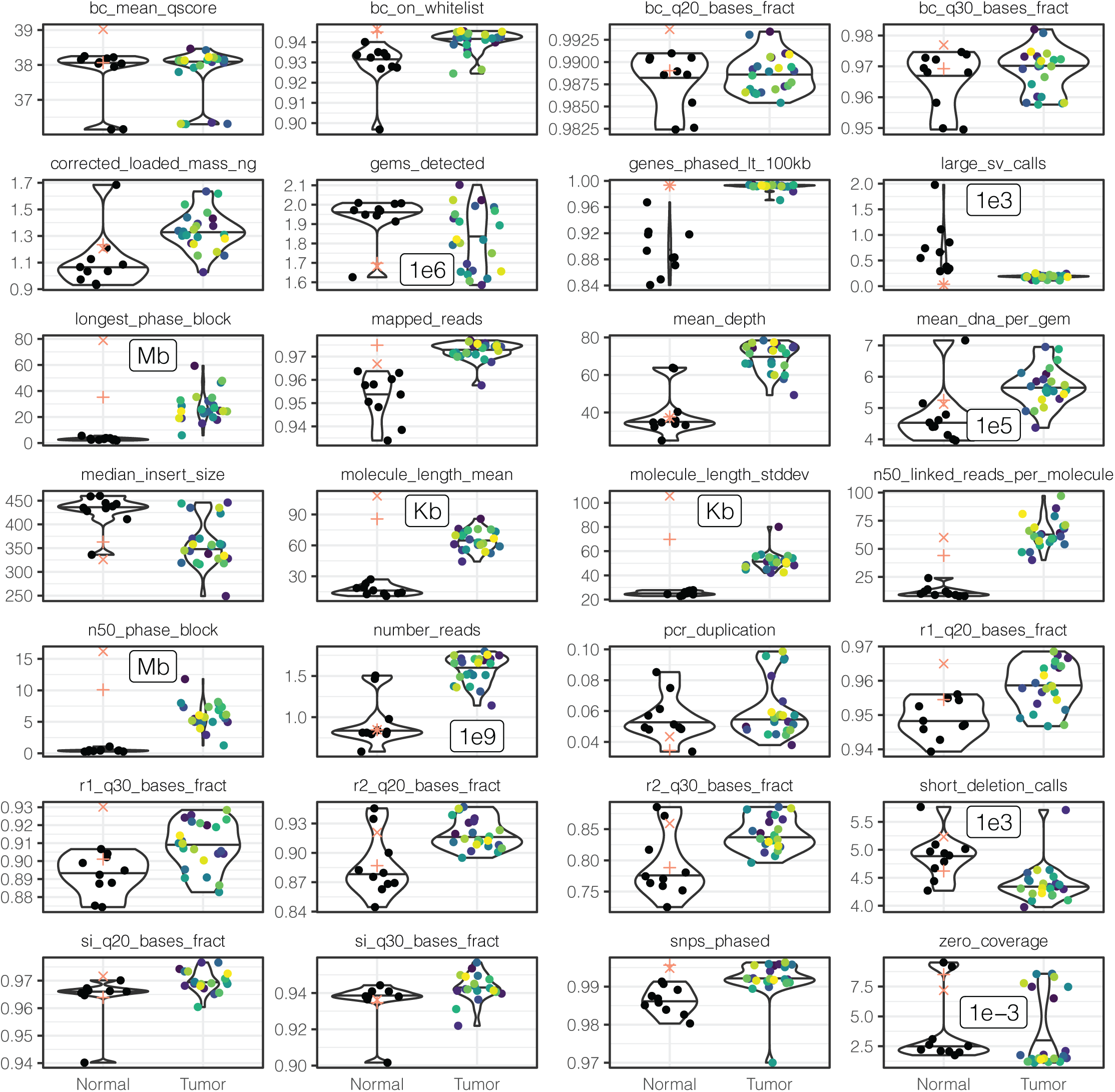
Phasing performance quality control summary measures for our tumor and normal samples plus 1000 Genomes samples NA12878 (+) and NA19240 (x). Violin plots defined as: center line, median; violin limits, minimum and maximum values; points, every observation. Definitions of metrics may be found here: https://support.10xgenomics.com/genome-exome/software/pipelines/latest/output/metrics.

**Supplementary Figure 2.**
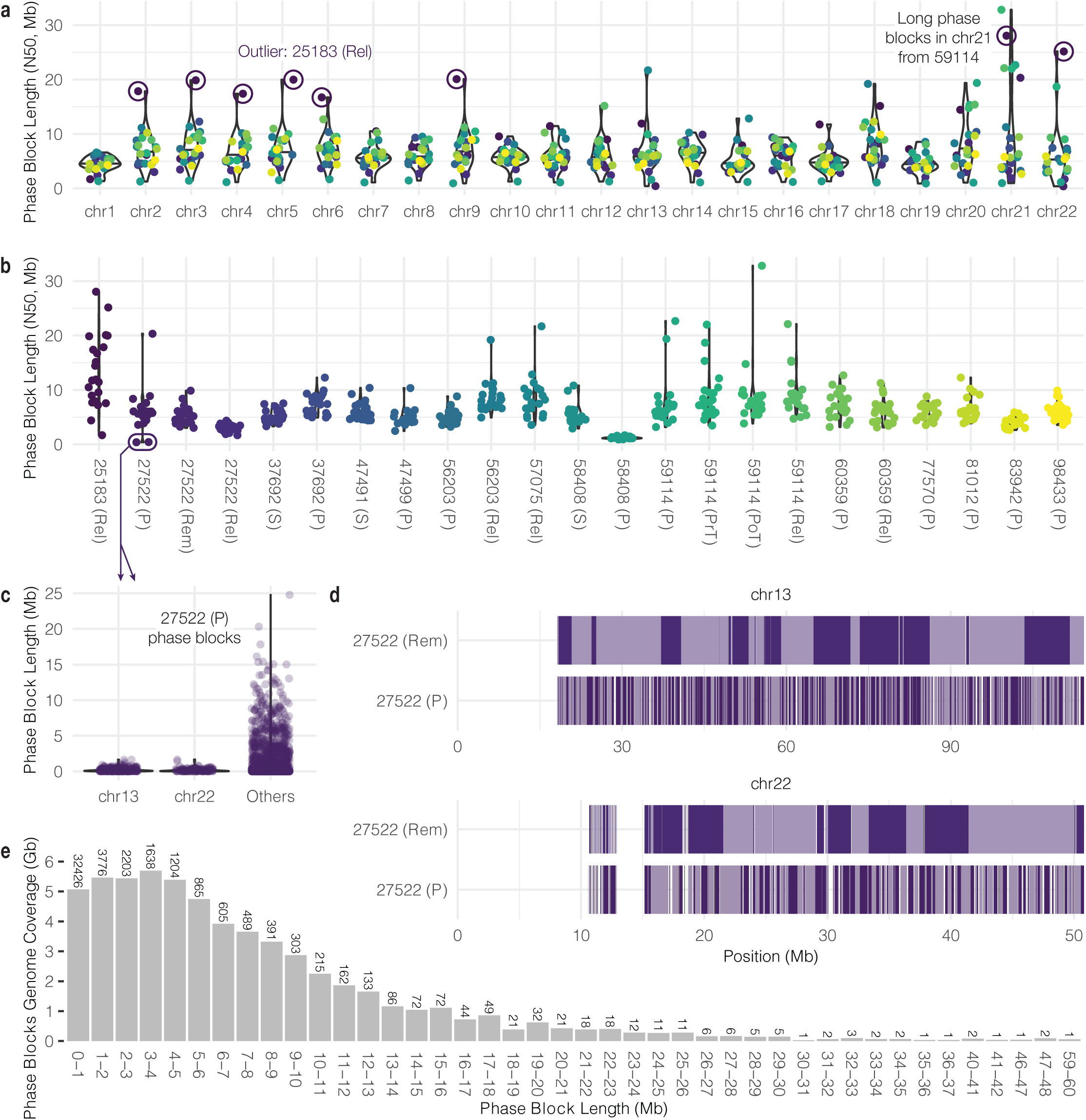
Phase block length distribution. **a.** Phase block length by chromosome across all samples. Outlier phase blocks from sample 25183 (Rel) circled. Violin plots defined as: center line, median; violin limits, minimum and maximum values; points, every observation. **b.** Phase block length per sample across all chromosomes. **c.** Phase block lengths of chr13, chr22, and others from 27522 (P). Phase blocks less than 1 kb filtered out for plotting. **d.** Chr13 and chr22 phase block boundaries from 27522 (P) and 27522 (Rem). Alternating dark and light boxes indicate adjacent phase blocks. **e.** Total phase block genome coverage from all samples combined, grouped by phase block length.

**Supplementary Figure 3.**
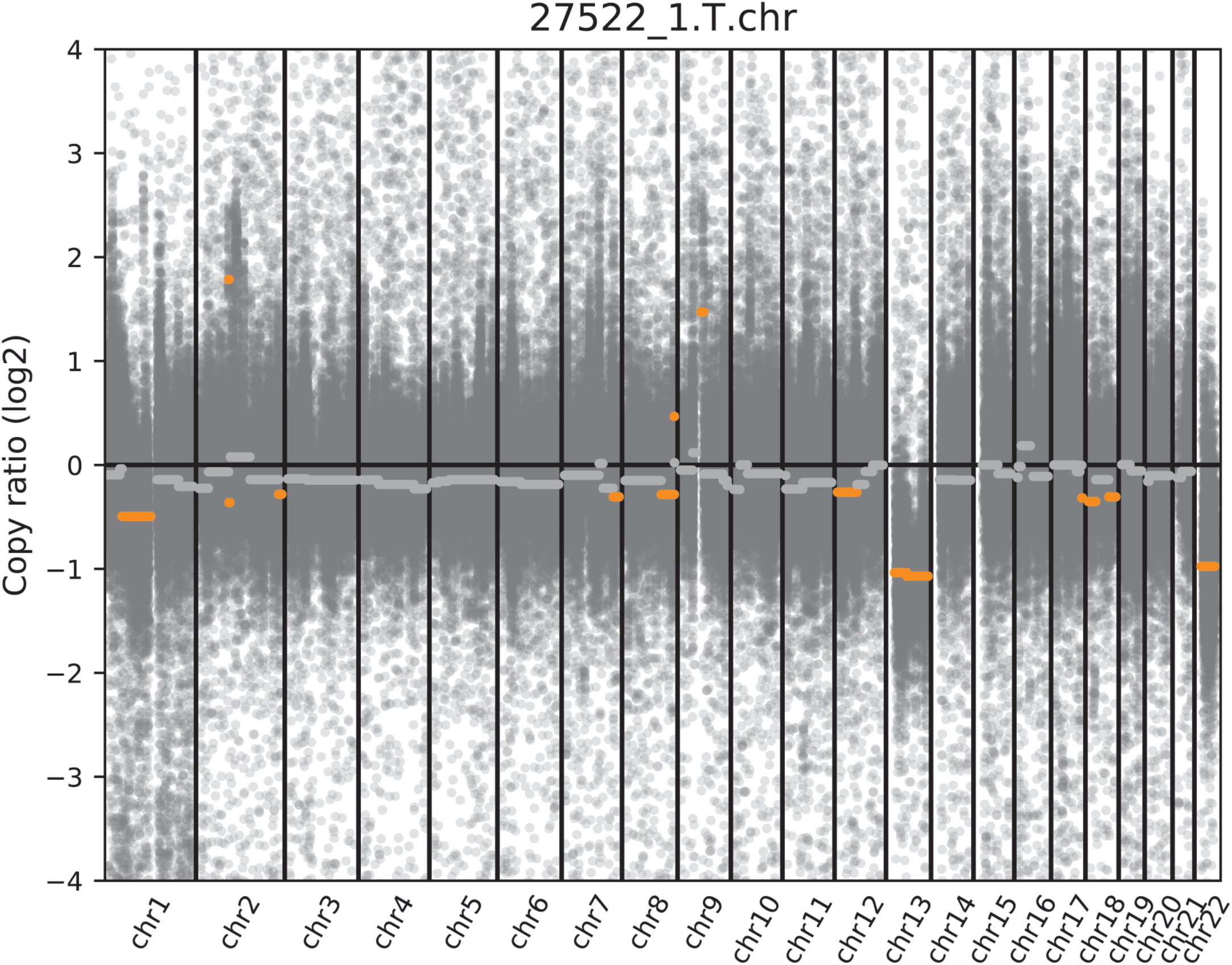
Copy number profile of Patient 27522 at the primary disease stage. Y-axis values are copy number ratios on the log2 scale.

**Supplementary Figure 4.**
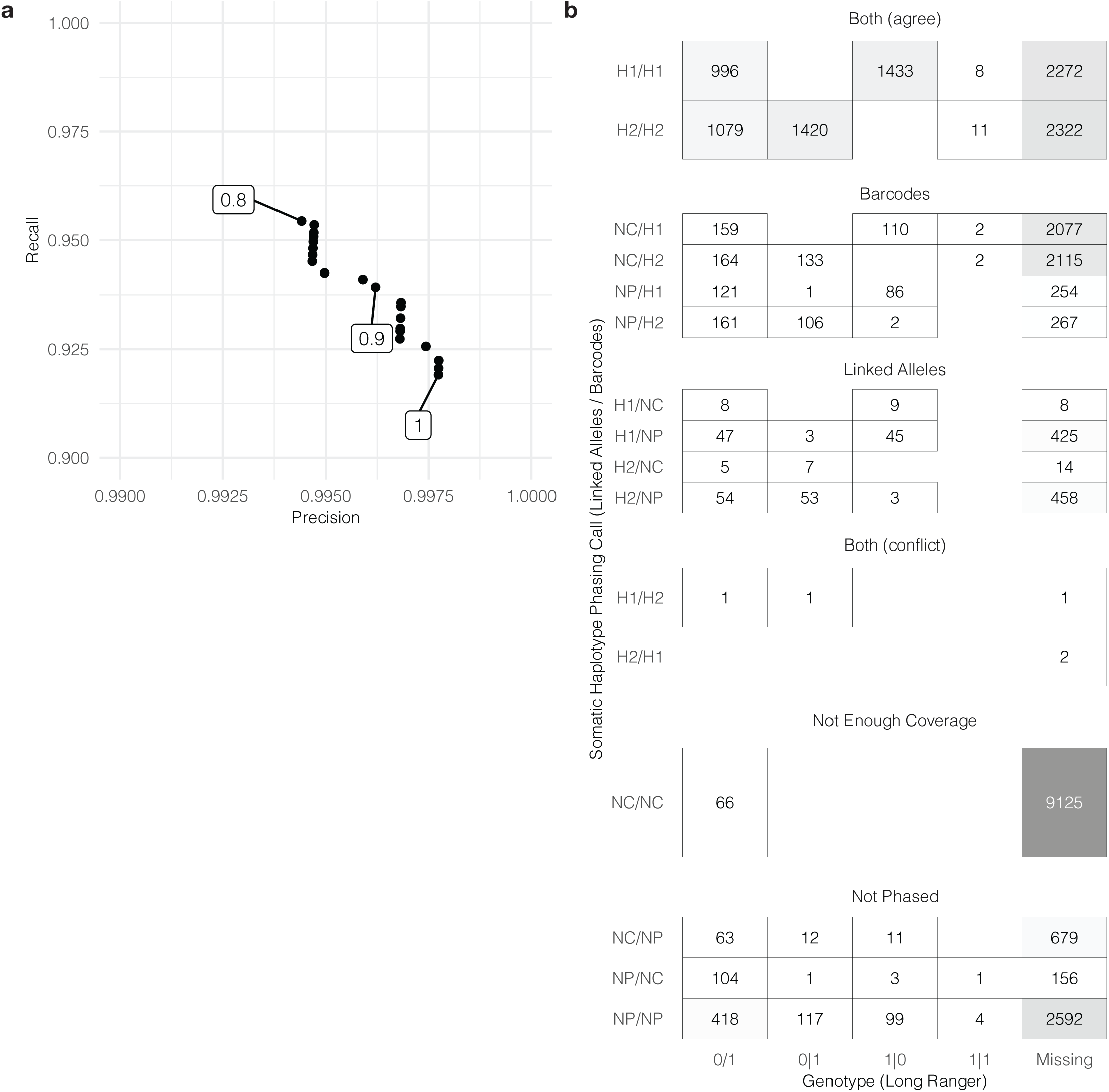
Additional information related to somatic mutation phasing. **a.** Precision/recall rates at various cutoffs for the proportion of linked-alleles assigned to one haplotype. **b.** Comparison of phasing results with Long Ranger genotypes.

**Supplementary Figure 5.**
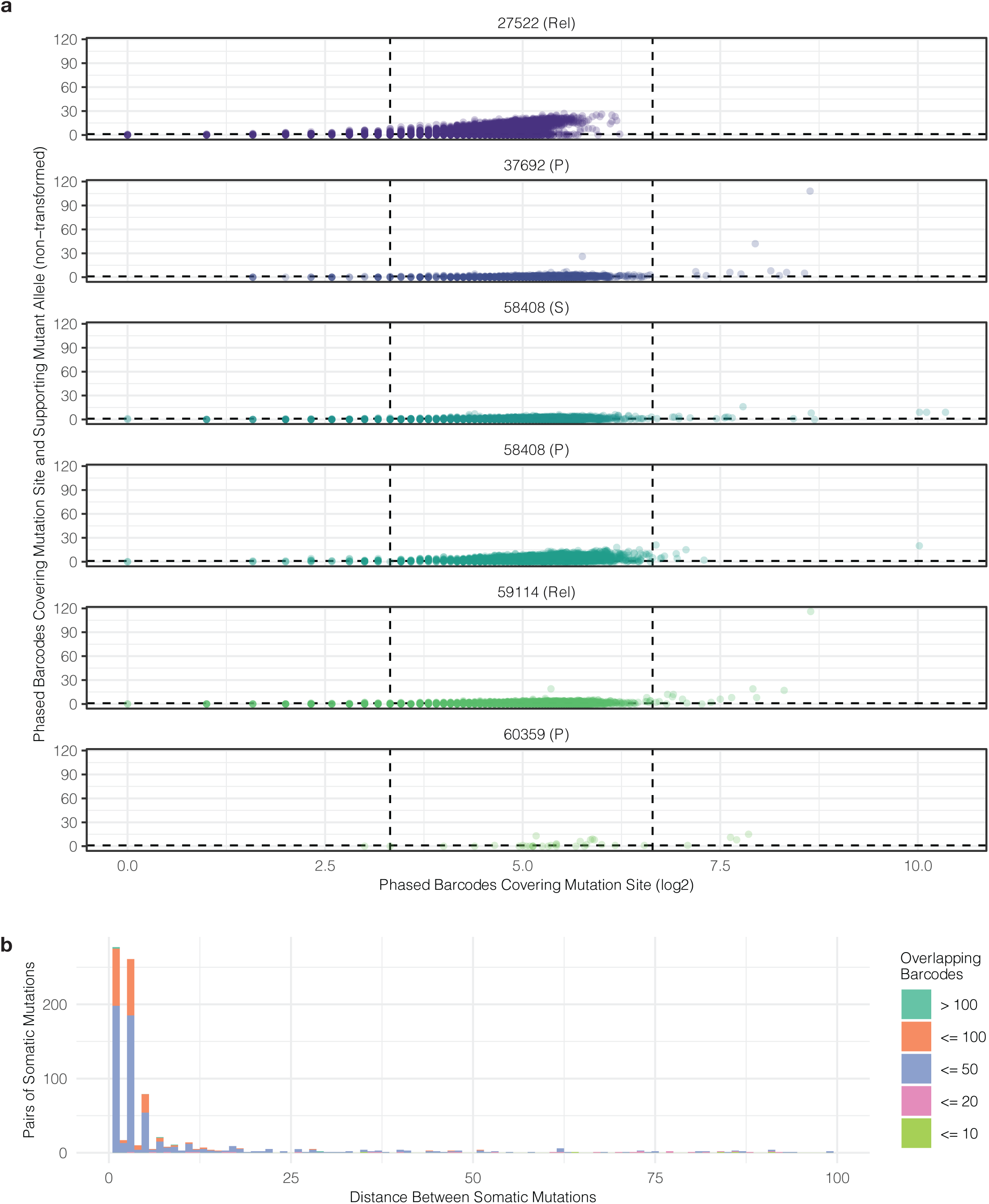
Additional information related to the relationship of pairs of somatic mutation. **a.** Number of barcodes covering each mutation site and those supporting the mutant allele. **b.** Number of overlapping barcodes by distance between somatic mutations less than 100 bp apart.

**Supplementary Figure 6.**
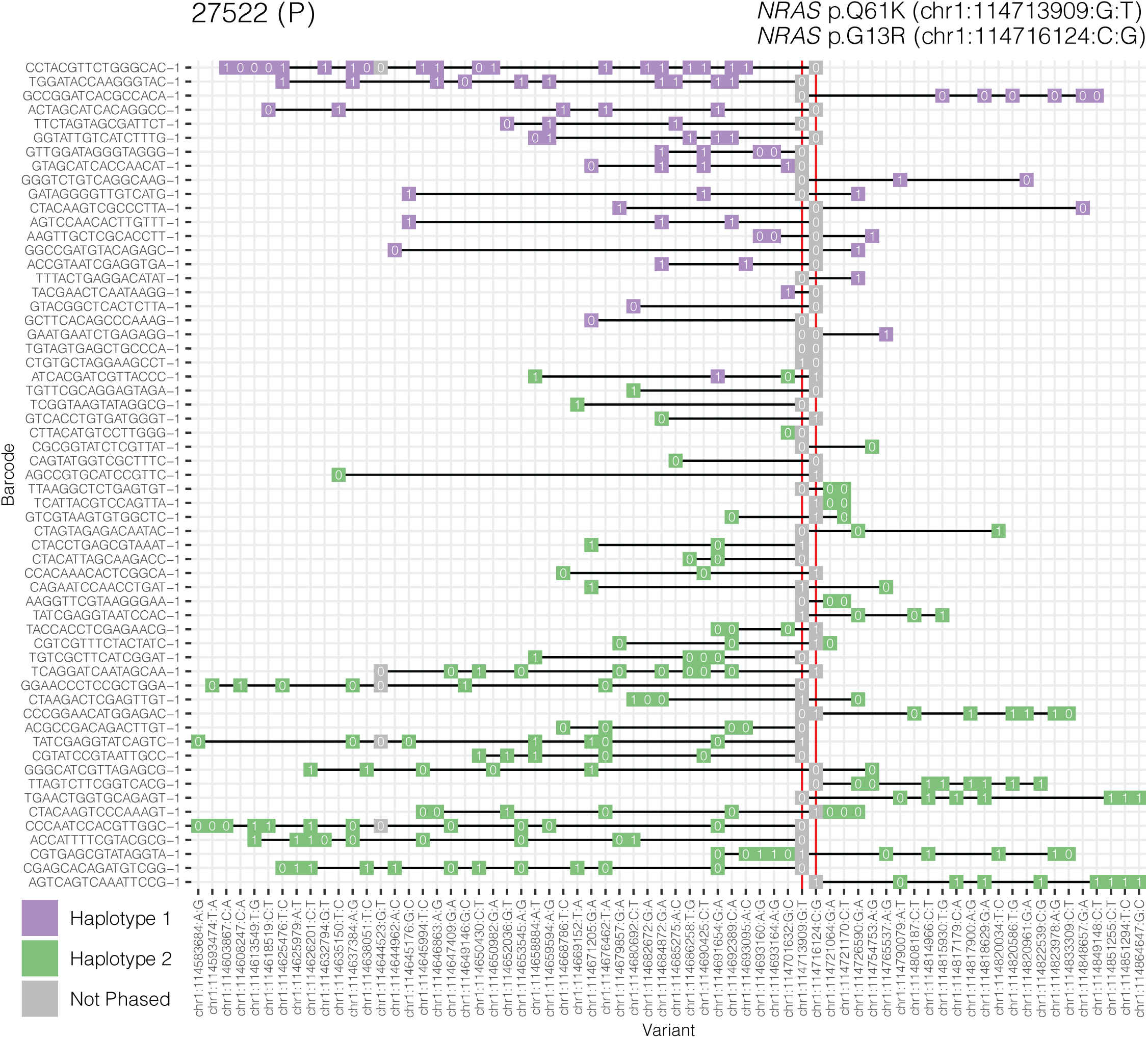
Barcodes supporting 27522 (P) NRAS hotspot mutation pair.

**Supplementary Figure 7.**
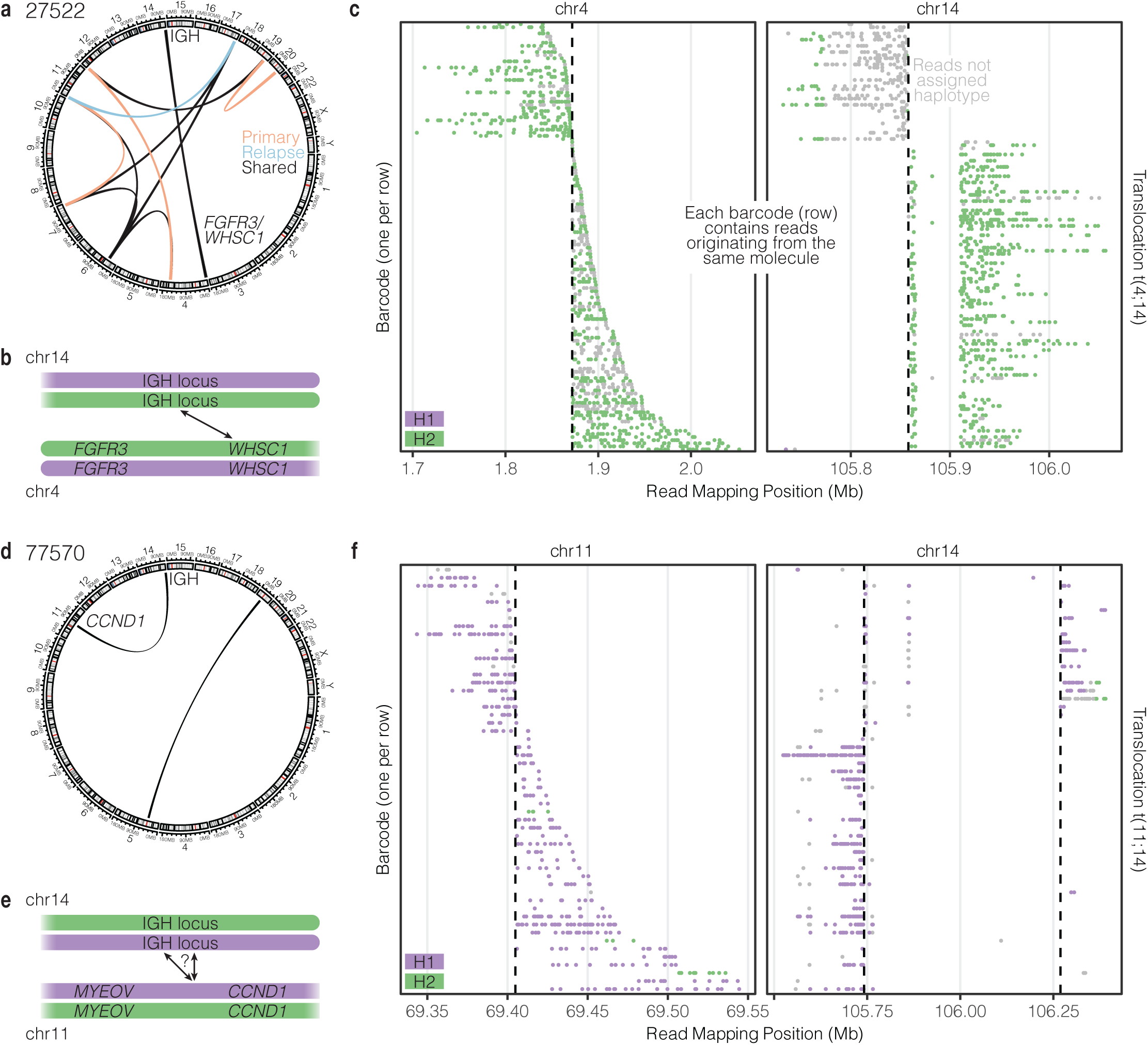
Common myeloma translocations mapped to haplotypes. **a.** Overlap of translocations observed in 27522 (P) and (Rel). **b.** Model of t(4;14) translocation. **c.** Barcodes supporting t(4;14) indicate a single haplotype origin. **d.** Translocations observed in 77570 (P). **e.** Model of t(11;14) translocation. **f.** Barcodes supporting t(11;14) indicate a single complex event.

**Supplementary Figure 8.**
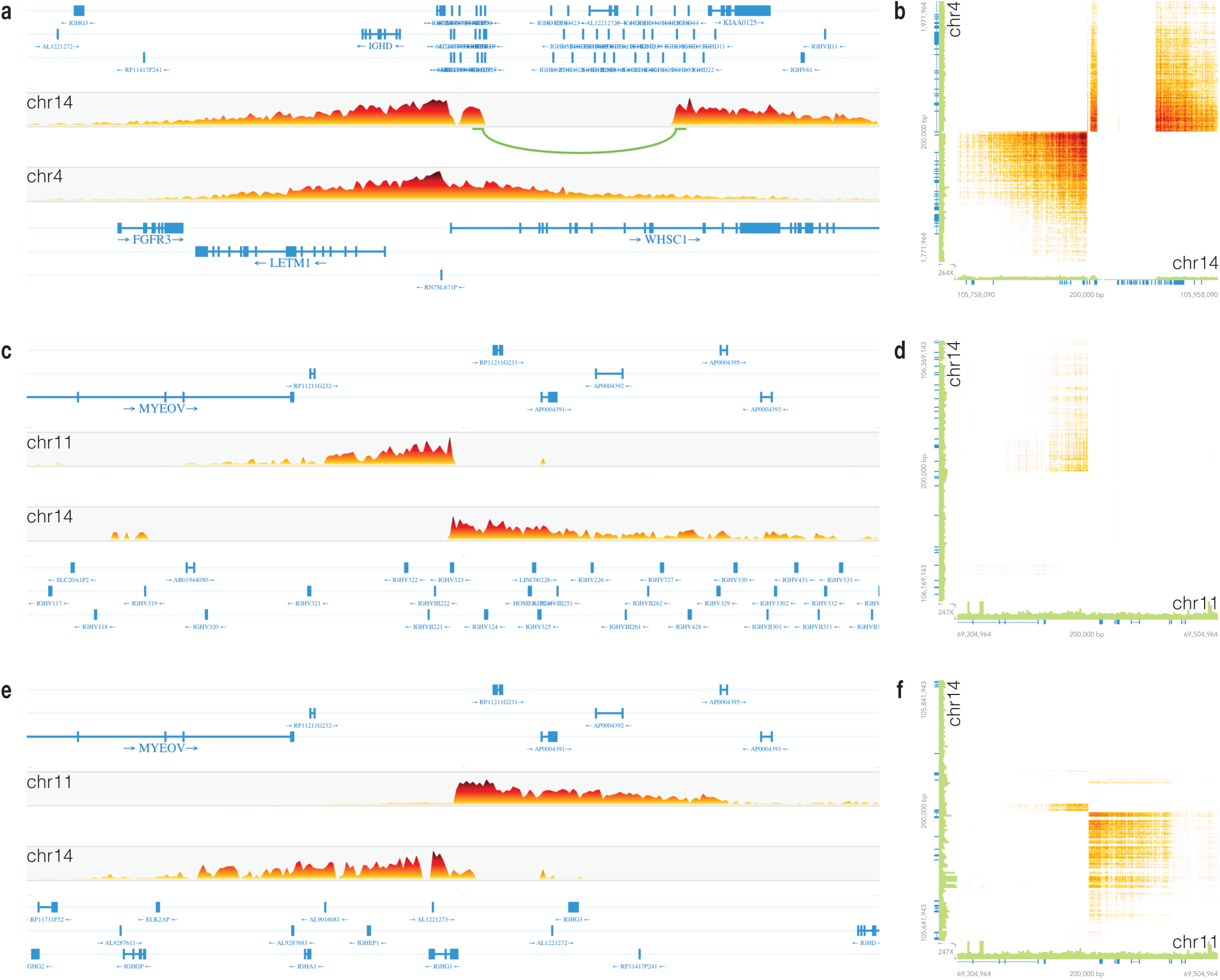
Barcode support for common myeloma translocations. **a-b.** 27522 (P) t(4;14). **c-f.** 77570 (P) t(11;14).

